# Chemoarchitectural influences on the cortical connectome confer resilience in aging

**DOI:** 10.64898/2026.04.23.719924

**Authors:** Santiago I. Flores-Alonso, Jack P. Solomon, Simon Dobri, Anthony R. McIntosh, Alex I. Wiesman

## Abstract

The macroscale signaling of the brain emerges from the integration of connected areas, orchestrated by microscale, regional molecular processes. Despite growing multimodal neuroimaging data resources, the role of cortical chemoarchitecture in shaping inter-regional functional connectivity remains poorly understood. Here, we examine whether brain regions that share a chemoarchitectural signature exhibit strong frequency-resolved functional connectivity, and whether aging moderates this neurochemical-functional alignment. Using magnetoencephalography data from across the healthy adult lifespan (n = 569), we identify a frequency-dependent organization of functional connectivity by slow neuromodulator systems, with low-frequency bands (*θ*-*α*) shaped most strongly by noradrenergic systems, and faster (*β*) alignment dependent on serotonergic chemoarchitecture. Aging strengthens the influence of neurochemistry on inter-regional connectivity, with frequency-specific implications for age-related cognitive performance. Neurochemical influences on *θ*-band connectivity were associated with worse cognition in older adults, while the opposite was true for the low-*γ* (*γ*_↓_) band. This suggests that neuromodulatory preservation of high-frequency dynamics in older adults may reflect neural resilience. Together, our findings indicate that neurochemical-functional alignment is frequency-dependent, distinguishing between maladaptive and resilient modes of brain organization.

## Introduction

The macroscale functional architecture of the human brain arises from the dynamic integration of distributed, reciprocally connected regions rather than from isolated regional computations alone (1; 2). These dynamics are rooted in region-specific molecular and cellular processes (3). Neurotransmitter systems regulate neuronal excitability, modulate firing probability, and tune synaptic gain, thereby governing the efficiency and fidelity of signal transmission. Through this modulation of cellular dynamics, neurochemical gradients effectively constrain the temporal coordination and frequency-specific communication that underlie network-level integration (4) and ultimately function (5). Importantly, cortical chemoarchitecture is itself systematically organized, mirroring core principles of structural and functional networks, such that molecular similarity between regions recapitulates canonical axes of cortical hierarchy and connectivity, implicating molecular heterogeneity as a substrate for functional flexibility (6; 7).

Critically, neurochemical systems do not exert uniform effects across temporal scales; rather, they differentially shape frequency-specific oscillatory processes that scaffold large-scale communication, which provides a temporal “clocking” mechanism to organize information transmission encoded as coordinated bursts of action potentials (8; 9). Neuromodulatory systems have been shown to exert frequency-specific influences on neural oscillations (10; 11), further supporting the idea that distinct neurochemical systems bias communication within specific spectral regimes. Within this framework, inter-regional synchronization at specific oscillatory frequencies operates as a dynamic gating mechanism, regulating when and where information is propagated, integrated, or suppressed across networks (12). Regional chemoarchitecture, in turn, likely constitutes a molecular substrate that biases the emergence and cross-regional coordination of these distinct oscillatory functional connectivity (FC) regimes.

Recent advances in positron emission tomography (PET) and open science have produced high-resolution atlases of neurotransmitter receptors and transporters, enabling a normative description of cortical chemoarchitecture that can be explicitly linked to systems-level brain organization (13). Work in this domain has progressed from demonstrating that neurotransmitter systems are dynamically coupled to whole-brain activity to formalizing chemoarchitecture as an additional organizational layer of the cortex (3). However, existing studies have focused almost exclusively on hemodynamic FC networks derived from functional magnetic resonance imaging (fMRI) (6; 7; 14; 15), and local patterns of frequency-defined cortical signaling from magnetoencephalography (MEG) (10; 16; 17; 18; 19; 20), leaving a critical gap in our understanding of how neurochemical architecture shapes band-limited oscillatory FC in the human brain.

The efficiency and flexibility of oscillatory FC networks are thought to support cognitive performance across the lifespan, with rhythmic synchronization enabling the selective routing and integration of information across distributed networks (21; 22). These effects are inherently band-limited, supporting distinct cognitive operations, including working memory, attention, memory retrieval, and executive control, through coordinated interactions among frontoparietal, frontotemporal, prefrontal, and thalamocortical systems (23; 24; 25; 26; 27; 28; 29; 30). Age-related changes in FC are heterogeneous, and substantial inter-individual variability suggests that preserved network organization may confer resilience against structural and molecular deterioration (31; 32). Contemporary frameworks of cognitive reserve emphasize the maintenance of efficient large-scale communication as a key determinant of successful aging (33). From this perspective, the degree to which large-scale communication is shaped by its underlying neurochemistry may represent a mechanistic substrate of reserve, reflecting the capacity of neuromodulator-constrained network dynamics to preserve efficient information routing despite aging-related perturbations to inter-regional morphometry (34), white matter (35), and metabolic integrity (36; 37).

Converging evidence suggests that this chemo-functional coupling is not only relevant for normative aging, but also reflects a key axis of vulnerability in neurodegenerative disease. Disruptions in the establishment and coordination of these frequency-specific communication channels have been linked to impaired cognition and are observed across neuropsychiatric and age-related neurodegenerative conditions, including notable aberrant network interactions in Parkinson’s (21; 38; 39; 40) and Alzheimer’s disease (23; 41; 42). Consistent with this, electrophysiological dynamics measured with MEG show robust alignment with underlying cellular and molecular organization (18; 43), implicating specific neurochemical systems in symptom heterogeneity in these conditions (19; 44). Together, these findings suggest that neurochemical-functional correspondence may provide a principled window into neurodegenerative vulnerability, linking molecular architecture to large-scale network dysfunction (45).

These integrative frameworks emphasize the need to reconcile connectivity blueprints spanning molecular and functional domains (46) as a critical dimension of brain dynamics in the face of aging. To close this gap, we tested whether normative neurotransmitter architecture aligns with frequency-resolved cortical FC and examined how this alignment relates to aging and cognitive function. We hypothesized that frequency-defined connectivity would be higher between regions with greater neurochemical similarity, reflecting a chemoarchitectural constraint on oscillatory cortical communication. We further hypothesized that this neurochemical–connectivity alignment would exhibit frequency-dependent variation. Finally, we posited that the strength of neurochemical shaping of FC would show frequency-specific associations with aging and cognitive abilities, such that strengthened alignment in select oscillatory frequencies indexes resilience-related mechanisms supporting cognitive reserve.

## Results

### Chemoarchitecture facilitates frequency-specific long-distance connectivity

To test whether similarity of cortical neurochemical profiles facilitates frequency-resolved functional connectivity, we integrated task-free MEG and cognitive data from the Cambridge Centre for Ageing and Neuroscience (Cam-CAN; n = 569, mean age=53.11 ± 17.4 years, 292 males) with PET-derived cortical atlases of 19 neurotransmitter receptor and transporter systems from neuromaps (Fig. 1). We first quantified correspondence between regional neurochemical similarity and band-limited FC across six canonical frequency ranges (*δ* : [1 − 4], *θ* : [4 − 7], *α* : [8 − 12], *β* : [15 − 29], *γ*_↓_ : [30 − 59], *γ*_↑_ : [60 − 90]*Hz*), adjusting for age, sex, and inter-regional distance. Linear mixed effects models revealed stronger connectivity between cortical regions with greater neurochemical similarity, and this effect was dependent on frequency band (Fig. 2A; *t* =[*δ* : 13.49, *θ* : 19.25, *α* : 26.59, *β* : 44.95, *γ*_↓_ : 24.32, *γ*_↑_ : −2.85]; *p*_*FDR*_([*δ, θ, α, β, γ*_↓_])*<* 0.001; *p*_*FDR*_(*γ*_↑_) *>* 0.05). Neurochemical-connectivity alignment increased from slow frequency *δ* and *θ* rhythms into the *α*- and *β*-bands, before decreasing markedly in the higher *γ* frequencies. Importantly, all reported effects remained significant after controlling for age-related structure in the neurochemical similarity matrix with p-values unchanged relative to the main analysis (*t* = [*δ* : 12.60, *θ* : 18.23, *α* : 24.68, *β* : 46.52, *γ*_↓_ : 24.54, *γ*_↑_ : −3.39]; *p*_*FDR*_ ([*δ, θ, α, β, γ*_↓_]) *<* 0.001; *p*_*FDR*_(*γ*_↑_) *>* 0.05). This indicates that the observed alignments cannot be attributed to non-specific age-related effects on PET tracer uptake (Fig. S1).

**Fig 1.**
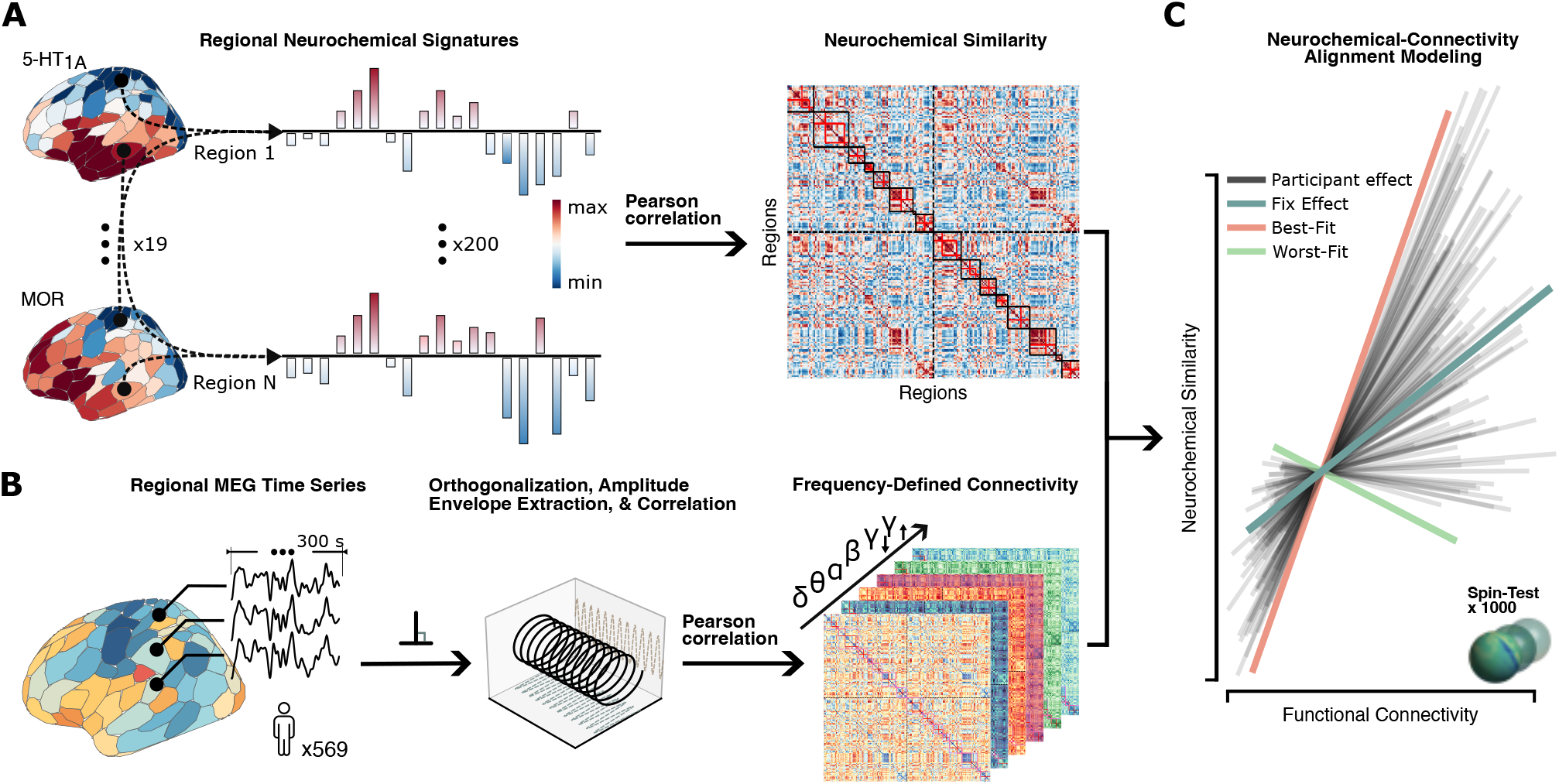
Quantification of alignment between neurochemistry and functional connectivity. Schematic overview of the multimodal pipeline integrating source-reconstructed MEG data with PET-derived neurochemical atlases. (**A**) Normative neurotransmitter receptor and transporter PET tracer maps were parcellated and used to compute pairwise neurochemical similarity via Pearson correlation. (**B**) Task-free MEG time series were filtered into 6 canonical frequency bands, parcellated, pairwise orthogonalized, Hilbert-transformed to extract the envelope amplitude, and used to derive frequency-resolved orthogonalized amplitude-envelope correlation (oAEC). (**C**) Linear mixed effects modeling was used to quantify the association between neurochemical similarity and oAEC, with participant-specific random slopes capturing individualized deviations from the population-level alignments.

**Fig 2.**
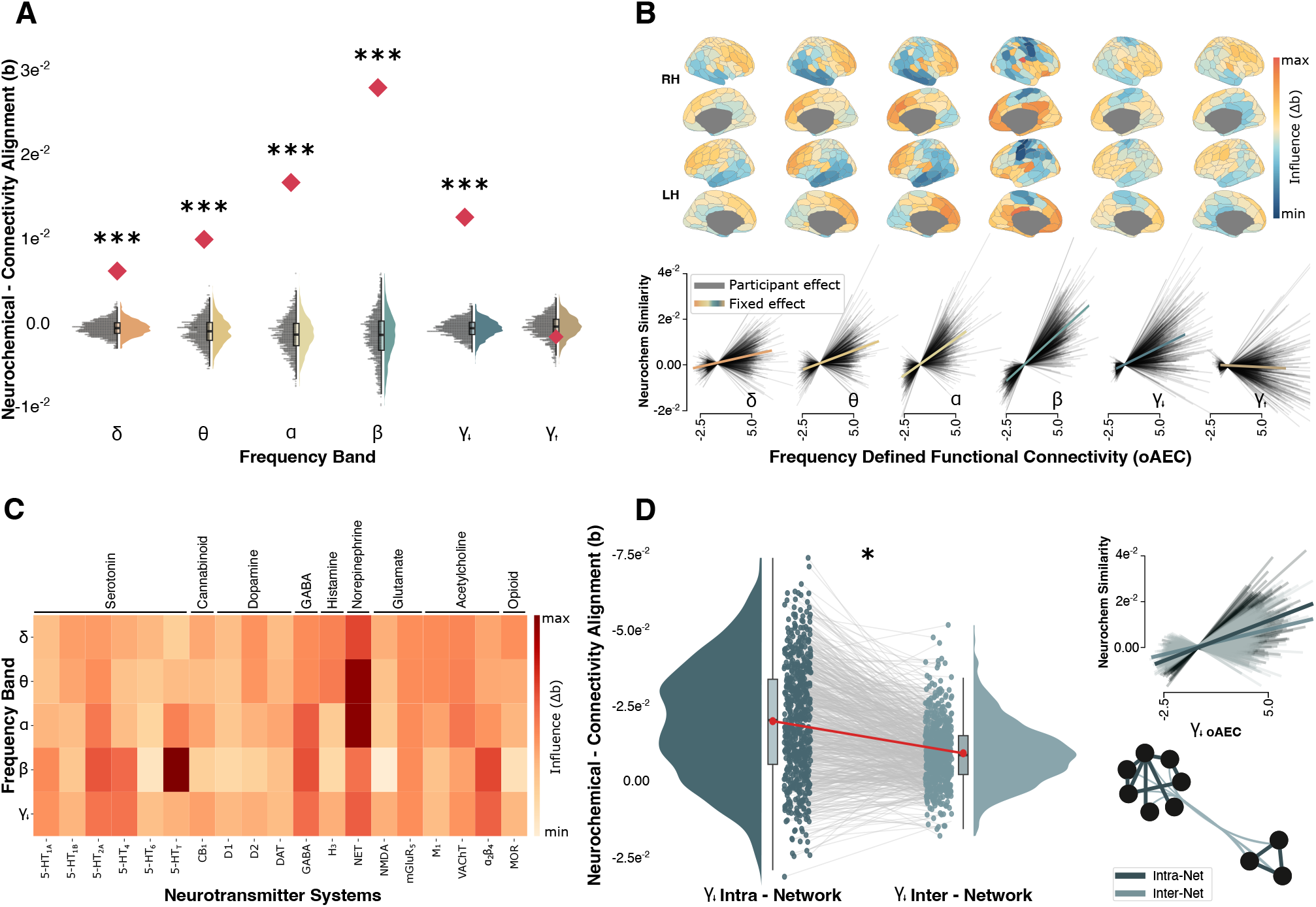
Frequency-resolved functional connectivity is shaped by neurochemistry. (**A**) Amplitude-envelope correlations (oAEC) and cortical chemoarchitecture show a robust frequency-dependent correspondence. Linear mixed effects models revealed positive neurochemical–connectivity alignment in δ, θ, α, β and γ_↓_-bands, with the largest effect in β. Red diamonds denote empirical fixed-effect estimates, and violins indicate permuted autocorrelation-preserving null distributions. (**B**) Brain maps show the frequency-specific change in model fit (Δb) after removing each ROI. Participant-level slopes illustrate how individual variability tracks the group-effect alignment across frequencies. (**C**) Neurotransmitter-specificity analyses demonstrate that alignment is differentially shaped by neuromodulatory systems. Serotonergic (5-HT) and noradrenergic (NET) systems exert the strongest influence across δ-β bands, whereas GABAergic, glutamatergic, cholinergic, and opioid systems exhibit weaker effects. (**D**) Participant-level alignments (dots) are higher when only considering intra-network connections, indicating that γ_↓_ neurochemical-functional coupling is primarily supported by within-network interactions. ∗p < 0.05, ∗ ∗ p < 0.01, ∗ ∗ ∗p < 0.001.

We next determined which cortical areas exert the strongest influence on these neurochemical–connectivity alignments using a regional leave-one-out (LOO) analysis in which each parcel was iteratively removed from the connectivity and neurochemical matrices, and the resulting change in the regression slope (Δ*b*) mapped onto the cortical surface. Across frequency bands, frontoparietal regions disproportionately drove neurochemical–connectivity alignment, while temporo-occipital and somatomotor parcels exerted minimal influence (Fig. 2B).

To identify which specific neurotransmitter systems most strongly influence functional connectivity, we performed a neurotransmitter atlas-level LOO analysis, where the neurochemical similarity matrix was recomputed with each neurochemical system removed, and the resulting change in neurochemical-connectivity model fit quantified (Fig. 2C). In the *θ*- and *α*-bands, the norepinephrine transporter (NET) exhibited the strongest influence on low-frequency neurochemical-functional alignment. In the *β*-band, alignments were most strongly driven by serotonergic systems, particularly the serotonin transporter (5-HTT). GABAergic and glutamatergic receptors exerted comparatively weaker influence on FC across frequency bands, likely due to their widespread but non-specific cortical expression. Finally, connectivity in the *δ* and *γ*_↓_-bands displayed minimal perturbation upon removal of individual neurotransmitter systems, suggesting that connectivity in these frequencies may be the result of multi-system neuromodulatory influences.

Next, we tested whether frequency-defined neurochemical alignment was preferentially supported by within-network or between-network interactions. Node-wise intra-/inter-network correspondence was defined according to the canonical resting-state cortical network organization proposed by Yeo et al. (47). Across the *δ*–*β* bands, neurochemical-functional alignment did not significantly differ within versus between networks (*t* = [*δ* : −1.11, *θ* : −23.55, *α* : −59.64, *β* : −68.65]; *p*_*FDR*_ *>* 0.05), indicating that the influence of neurotransmitter systems on neurophysiological signaling at these frequencies is not preferentially driven by either intra- or inter-network connectivity. In the *γ*_↓_-band, however, intra-network neurochemical-connectivity alignments were stronger than inter-network alignments (*t* = [*γ*_↓_ : −52.96]; *p*_*FDR*_ = 0.031; Fig. 2D).

### Increased neurochemical-connectivity alignment in aging confers cognitive resilience

Finally, we examined whether the strength of neurochemical–connectivity alignments differed as a function of aging and age-related cognitive variations (Fig. 3A). We focused on cognitive scores from the Cattell Culture Fair Intelligence Test, given the preferential age-related declines in fluid versus crystallized intelligence (48; 49). The *θ, β* and *γ*_↓_-bands showed an age-related modulation of neurochemical-functional alignment (*t* = [*δ* : 0.91, *θ* : 1.23, *α* : 0.98, *β* : 8.30, *γ*_↓_ : 1.51; *p*_*FDR*_ = [*δ* : 0.37, *θ* : 0.049, *α* : 0.079, *β* : 0.0006, *γ*_↓_ : 0.048]; Fig. 3B). With the exception of the *β*-band (*t* = [*δ* : 0.28, *θ* : −0.38, *α* : 0.35, *β* : 0.78, *γ*_↓_ : 0.91]; *p*_*FDR*_ = [*δ* : 0.114, *θ* : 0.010, *α* : 0.615, *β* : 0.114, *γ*_↓_ : 0.006]; Fig. 3B),neurochemical-connectivity alignment in these frequencies was also associated with age-corrected fluid intelligence performance. In the *γ*_↓_ band, individuals expressing stronger-than-expected neurochemical-connectivity alignment for their age exhibited preserved fluid intelligence (*r* = .11; *p*_*FDR*_ = 0.006), indicating a resilient or neuroprotective effect of neurochemical organization of FC (Fig. 3C). Neurochemical-connectivity alignment in the *θ*-band exhibited an opposite pattern (*r* = −0.09; *p*_*FDR*_ = 0.01).

**Fig 3.**
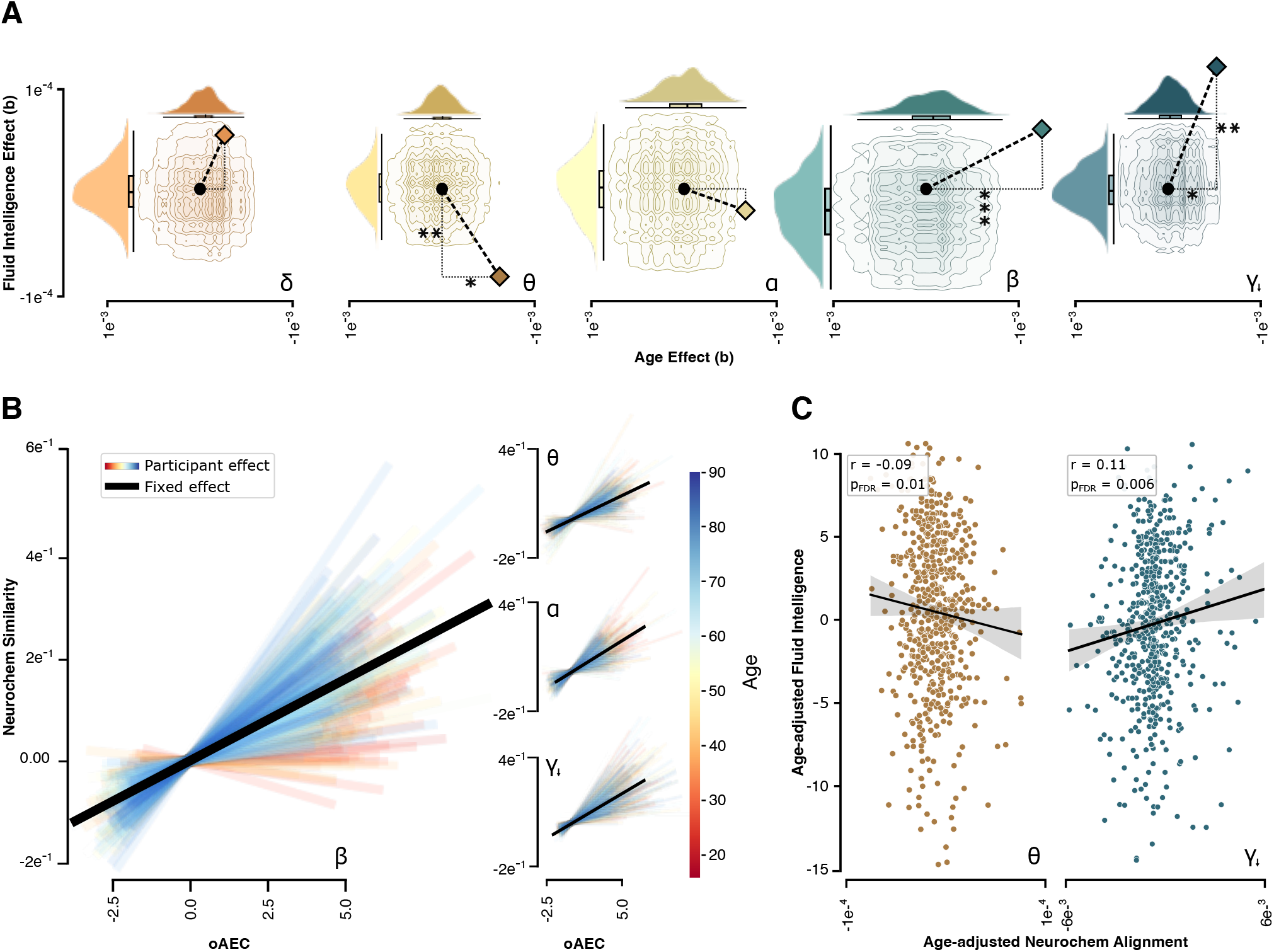
Aging and cognition modulate frequency-resolved neurochemical–connectivity alignment. (**A**) Null distributions (displayed as both joint-topographic and independent density plots) for the interaction between neurochemical alignment and age (x-axis) and age-adjusted fluid intelligence (y-axis) for each frequency band. Diamonds indicate the empirical model estimates for the joint effect; dotted lines indicate their deviations from the null centroid; asterisks denote significant departures along each dimension. (**B**) Group-effect (black) and participant-level (colored) slopes indicate the age effects at the individual level. (**C**) Age-adjusted associations between neurochemical alignment and fluid intelligence for θ and γ_↓_. ∗p < 0.05, ∗ ∗ p < 0.01, ∗ ∗ ∗p < 0.001.

## Discussion

By integrating MEG-derived FC with a multi-system atlas of inter-regional neurochemical similarity, we demonstrate that large-scale communication is related to the brain’s underlying neurochemical landscape. This neurochemical-functional alignment is both frequency- and neurotransmitter-dependent, indicating that distinct neuromodulatory systems selectively shape oscillatory communication regimes. Aging strengthened the influence of neurochemical systems on functional connectivity, and individuals with greater high-frequency neuromodulatory influences on connectivity exhibited preserved fluid intelligence relative to age-matched peers. Together, these findings position cortical neurochemistry as a potential mechanistic scaffold linking molecular architecture to frequency-specific communication and cognitive resilience in aging.

Frontoparietal regions had a pronounced influence on neurochemical-functional alignment across all frequency bands, which is likely due to the dense innervation of frontoparietal control networks by neuromodulator systems (50; 51; 52). The importance of these neurochemically-orchestrated network interactions for preserved cognition in aging is also intuitive, given the reliance of fluid intelligence on frontoparietal signaling (53). Leave-one-out analyses of each neurochemical system suggested that noradrenergic systems scaffold synchronization within low-to-intermediate frequency ranges, with serotonin playing an outsized role in connectivity at higher frequencies, positioning norepinephrine and serotonin as frequency-specific regulators of long-range oscillatory integration along these frontoparietal axes. Converging evidence supports these interpretations. Signaling in the *α*-band has been clearly linked to noradrenergic tone (54; 20), while genetic disruption of serotonin transporters reduces *β*-band FC (55) and pharmacological modulation via serotonergic psychedelics (e.g., LSD and psilocybin) alters *β*-band dynamics and connectivity (56; 57). In contrast, shaping of *γ*_↓_ connectivity by neurochemistry was not specific to a single neuromodulator and was also markedly stronger in within-network than between-network connections. This pattern is consistent with canonical theories of high-frequency oscillations, which are characterized instead by short-range connectivity governed by local microcircuit architecture and excitation–inhibition balance below the spatial resolution of MEG (9; 58). Collectively, these results support a frequency-resolved framework in which distinct neuromodulatory systems differentially structure large-scale versus local communication regimes.

Older adults exhibited stronger effects of neurochemistry on connectivity across multiple frequency bands. The strongest such aging effect emerged in the *β*-band, indicating that FC in this regime is increasingly shaped by neurochemistry as we age. However, the strength of this *β* alignment was not associated with cognitive performance. In contrast, similar neurochemical-connectivity effects in the *θ* and *γ*_↓_-bands were qualitatively less strongly related to age, but were significantly associated with cognition. Stronger *θ*-band alignment predicted poorer fluid intelligence, while stronger *γ*_↓_ alignment was associated with better cognitive performance, suggesting that preserved neurochemical-functional coupling at faster timescales may reflect more efficient or resilient local network dynamics. These findings are consistent with a neural slowing framework in which dominance of low-frequencies and impaired high-frequency signaling reflects reduced processing efficiency and altered excitation–inhibition balance (21; 50). This interpretation aligns with evidence from neurodegenerative diseases, where increased slow-frequency activity and spectral shifts are linked to amyloid and tau pathology, as well as cognitive decline (59; 60; 61).

The limitations of our study should be noted. Our analyses rely on normative neurochemical maps, which do not capture inter-individual variability. Although the spatial organization of neurochemical similarity appears highly stable across individuals (62; 63), subject-specific inference is not possible with these atlases, limiting interpretations regarding e.g., whether the effects we observe are due to variations in receptor and transporter densities or function. We are also unable to model age-related changes in integrity per each neurochemical atlas using this approach. This concern is partially mitigated by the fact that the underlying maps overlap entirely with the age range of the neuromaps participant sample (35–55 years), and by control analyses indicating that controlling for age effects on atlas-nonspecific tracer uptake yields near-identical effects (Fig. S1). Nonetheless, the gold standard approach remains participant-specific PET maps. Guided by our findings, future work should prioritize individual-level imaging of noradrenergic and serotonergic systems to provide nuance. We also anticipate that future research will extend these findings using longitudinal designs to resolve the temporal dynamics of neurochemical-functional alignments and their importance for cognitive reserve.

Together, our findings indicate that the shaping of FC by neurochemical systems is highly frequency- and age-dependent. We emphasize the importance of slow neuromodulator systems in inter-regional signaling, and that future research should consider frequency content in analyses of neurochemical influences on inter-regional connectivity regimes. We also find that these influences increase as we age, with neuromodulation of inter-regional *γ*_↓_ signaling indexing cognitive resilience. Clear next steps for this work include investigations into how neurodegenerative pathologies moderate these neurochemical-functional relationships, and whether these frequency-specific associations can be targeted by neurostimulation and pharmacological interventions to preserve cognition in individuals at high risk for these disorders.

## Methods

### Participants and data

Task-free MEG and structural MRI data were obtained from the Cambridge Centre for Aging and Neuroscience (Cam-CAN) repository, a cross-sectional cohort spanning the adult lifespan (64). Task-free MEG recordings were acquired using a VectorView system (Elekta Neuromag Oy, Helsinki) comprising 306 sensors (204 planar gradiometers, 102 magnetometers). Continuous head position was tracked via four head-position indicator (HPI) coils, enabling offline motion correction. Data were sampled at 1,000 Hz with an online anti-aliasing band-pass filter (0.03–330 Hz) and processed using MaxFilter for environmental noise suppression. Vertical and horizontal electrooculogram (EOG) signals were recorded using bipolar electrode pairs to monitor eye blinks and saccades, and an additional bipolar pair recorded the electrocardiogram (ECG) for identification of cardiac artifacts.

From the 647 participants available in the Cam-CAN dataset, we analyzed eyes-closed task-free MEG data from 569 neurologically healthy adults (age range = [18-90]; mean age = 53.11 ± 17.4 years; 292 males). Participants were excluded if MEG or structural MRI data failed preprocessing or source reconstruction quality control, or if source estimates exhibited abnormal cortical power topographies inconsistent with canonical frequency-band distributions (65). Each participant completed approximately 8 min 40 s of acquisition, with the first 20 s discarded to minimize transient adaptation effects. High-resolution structural MRIs were collected using a 3D T1-weighted MPRAGE sequence (TR = 2250 ms; TE = 2.99 ms; TI = 900 ms; flip angle = 9^*◦*^; FOV = 256 × 240 × 192 mm; voxel size = 1 mm isotropic; GRAPPA factor = 2; acquisition time = 4 min 32 s). Each individual’s T1-weighted MRI data were automatically segmented and labeled with Freesurfer (7.4.1) (66) recon-all. Noise statistics were derived from five-minute empty-room recordings acquired as close as possible in time to each participant’s MEG session.

Fluid cognitive performance was assessed using the Cattell Culture Fair Intelligence Test (CFIT) (67; 68), administered as part of the Cam-CAN neuropsychological battery. The CFIT is a standardized, nonverbal measure of fluid intelligence that evaluates abstract reasoning through matrix completion, classification, and series-based problem-solving tasks. Raw total scores were extracted from the repository and treated as a continuous variable reflecting fluid reasoning ability. To control for age-related effects, scores were residualized with respect to age using a linear model, such that the resulting values reflect age-independent variability in cognitive performance. Specifically, we estimated:

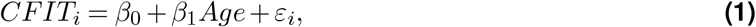

and defined the age-adjusted scores as the residuals:

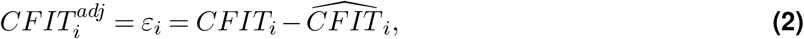

Where 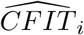 denotes the predicted score from the age model. These residualized scores were subsequently used in all analyses to isolate variance in fluid intelligence not attributable to normative aging effects.

The Cam-CAN study and open data release were approved by the local institutional review board, and all participants provided written informed consent. Secondary analyses of these data were also approved by the research ethics board at Simon Fraser University. All research protocols complied with the Declaration of Helsinki.

### MEG data processing

Task-free and empty-room MEG data were preprocessed in MNE-Python 1.10.2 (Python 3.13.9) following good-practice guidelines (69). Spatiotemporal signal-space separation (tSSS (70); window duration = 10 s, correlation limit = 99%) was applied to task-free data and empty-room recordings. Data were band-pass filtered (1–90 Hz), notch-filtered at 50 Hz, downsampled to 500 Hz, and cropped to a fixed interval across participants (30–330 seconds). We used electro-cardiogram and -oculogram recordings to define signal-space projections (71) (SSPs) for the removal of cardiac and ocular artifacts.

Coregistration between MEG sensor space and individual T1-weighted MRI anatomy was performed using the three fiducial landmarks (nasion and bilateral preauricular points) and 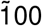 digitized head surface points. Given that the T1-weighted images were defaced–limiting the reliability of surface-based alignment, we instead used precomputed linear MRI–MEG transformation matrices (72) (https://github.com/tbardouille/camcan_coreg), ensuring consistent and robust cross-modal registration across participants. Forward solutions were computed using a three-layer Boundary Element Method (BEM) head model (73). Cortical source activity was reconstructed using linearly constrained minimum-variance (LCMV) beamforming (74). Data covariance matrices were estimated from the preprocessed sensor-level data and regularized (*λ* = 5%) to stabilize spatial filter estimation. Beamformer weights were computed across 20,484 cortical vertices per participant, with source orientations optimized in the direction of maximal projected power. Spatial filters were normalized using a unit-noise-gain constraint to reduce depth bias and ensure comparable sensitivity across cortical locations.

Cortical source time series were grouped into 200 regions of interest (ROIs) using the Yeo 200–17 network parcellation and reduced to a single representative signal per region via the first principal component. Frequency-resolved functional connectomes between all 200 ROIs were computed using orthogonalized amplitude-envelope correlation (oAEC (75)), yielding a symmetric 200 × 200 connectivity matrix per participant and frequency (*δ* : [1 − 4], *θ* : [4 − 7], *α* : [8 − 12], *β* : [15 − 29], *γ*_↓_[30 − 59], *γ*_↑_ : [60 − 90] Hz).

### Neurochemical similarity computation

Pairwise neurochemical similarity between ROIs was quantified using PET-derived cortical atlases of 19 neurotransmitter receptor and transporter systems obtained from neuromaps (13), parcellated into the same 200 cortical ROIs as the MEG data, producing a 200 × 19 matrix of cortical chemoarchitecture. For some neurotransmitter systems *ψ*, several PET tracers or participant cohorts contributed distinct maps. To place them on a common scale while accounting for sample size, each PET map *i* was z-scored across parcels and combined using a sample-size–weighted average:

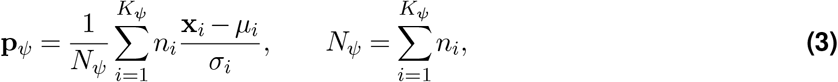

where **x**_*i*_ ∈ ℝ^200^ is the receptor/transporter density vector for the PET map *i, µ*_*i*_ and *σ*_*i*_ are the mean and standard deviation of **x**_*i*_, *n*_*i*_ is the sample size of the corresponding PET map, and *K*_*ψ*_ is the total number of maps available for the *ψ* system. The resulting **p**_*ψ*_ ∈ ℝ^200^ constitutes a z-scored and sample size-weighted PET atlas for the relevant neurotransmitter system.

The resulting 19 **p**_*ψ*_ vectors were stacked into a matrix **P** ∈ ℝ^200*×*19^ that represents the multi-receptor neurochemical profile of all ROIs. The pairwise neurochemical similarity between regions *i* and *j* was quantified as the Pearson correlation between their corresponding rows of this matrix.

### Neurochemical-connectivity alignment modeling

Neurochemical–connectivity alignment was assessed using a linear mixed effects model operating on vectorized connectivity and neurochemical similarity matrices. Model estimation used the *lme4*(76) package in R and maximum likelihood with the “*bobyqa*” optimizer. The lower triangular elements of the neurochemical similarity matrix were vectorized to form a single outcome vector *y* ∈ ℝ^*E*^ for these models, where *E* denotes the number of unique inter-regional edges. Predictor variables were constructed analogously by vectorizing the lower triangles of the frequency-resolved MEG FC matrices, yielding edge-wise connectivity vectors *x*_*pb*_ ∈ ℝ^*E*^ for each participant and frequency band. Participant-level demographics (i.e. age and sex) and Euclidean distance between ROIs centroids were included as covariates in all models. Random slopes and intercepts were specified at the participant level to capture inter-individual variability in the strength of neurochemical–connectivity alignment.

Formally, the model was specified as:

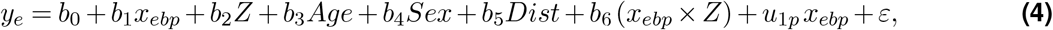

where *e* indexes edges (unique parcel pairs). *b*_0_ is the global intercept, and *b*_1_ captures the group-level relationship between frequency-specific connectivity and neurochemical similarity. Fixed effects *b*_2_–*b*_5_ adjust for moderator-specific effects (*Z*), demographic covariates (Age, Sex), and spatial constraints imposed by inter-regional distance (Dist). The interaction term *b*_6_ quantifies statistical moderation of the connectivity–neurochemical alignment by *Z*.

Random effects were specified as participant-specific slopes on connectivity: *u*_1*p*_ ∼ N (0, *τ* ^2^) captures participant-specific variability in the strength of this association. Residual error is denoted by *ε* ∼ N (0, *σ*^2^).Associations between neurochemical–connectivity alignment and participant-level measures (i.e., age, node-wise intra-/inter-network correspondence, and cognitive performance) were operationalized as moderation effects on the connectivity term via inclusion of the corresponding interaction *x*_*ebp*_ × *Z*.When age was the moderator (*Z* = Age), the term *b*_3_ Age was omitted to avoid redundancy.

### Null models and statistical significance

Statistical significance of neurochemical-connectivity alignments was assessed using spatial autocorrelation-preserving permutation tests (i.e., spin tests (77; 78)). Following the neuromaps framework, we generated a surface-based representation of the 200-parcel atlas on the FreeSurfer-fsaverage template using files from the Connectome Mapper Toolkit (https://github.com/LTS5/cmp). Each parcel was assigned spatial coordinates by selecting the spherical projection coordinates of the vertex closest to its center of mass. To generate spatially constrained null models, these parcel coordinates were rotated on the sphere, and each original parcel was reassigned the value of its nearest rotated neighbor (Hungarian method (79); 1,000 repetitions). Parcels mapping onto the medial wall were instead assigned the value of the next most proximal valid parcel. All rotations were performed at the parcel level, independently for each hemisphere, to avoid data upsampling and preserve hemispheric topology. Empirical estimates of frequency-specific neurochemical–connectivity alignment were compared against their corresponding spin-based null distributions using a two-tailed test, yielding nonparametric p-values that account for cortical autocorrelation, which were then adjusted across multiple comparisons to control the false discovery rate using the Benjamini-Hochberg procedure (80).

## ACKNOWLEDGEMENTS

This work was supported by the Canada Research Chair (CRC-2023-00300) in Neurophysiology of Aging and Neurodegeneration, a Discovery Grant from the Natural Sciences and Engineering Research Council of Canada (RGPIN-2025-04783), and infrastructure grants from the Canadian Foundation for Innovation and the British Columbia Knowledge Development Fund to AIW. Computing resources were provided in part by the Digital Research Alliance of Canada.

## DATA AVAILABILITY

The Cam-CAN dataset analyzed in this study is publicly available through the Cam-CAN data portal (https://camcan-archive.mrc-cbu.cam.ac.uk/). Neurochemical maps were obtained from PET-derived atlases available via neuromaps (https://github.com/netneurolab/neuromaps). Code used for MEG preprocessing is available via the McIntosh Lab GitHub repository (https://github.com/McIntosh-Lab/tvb-ccmeg), and analysis scripts developed for this study are available from the corresponding authors upon request.

## Supplementary Information

**Fig S1.**
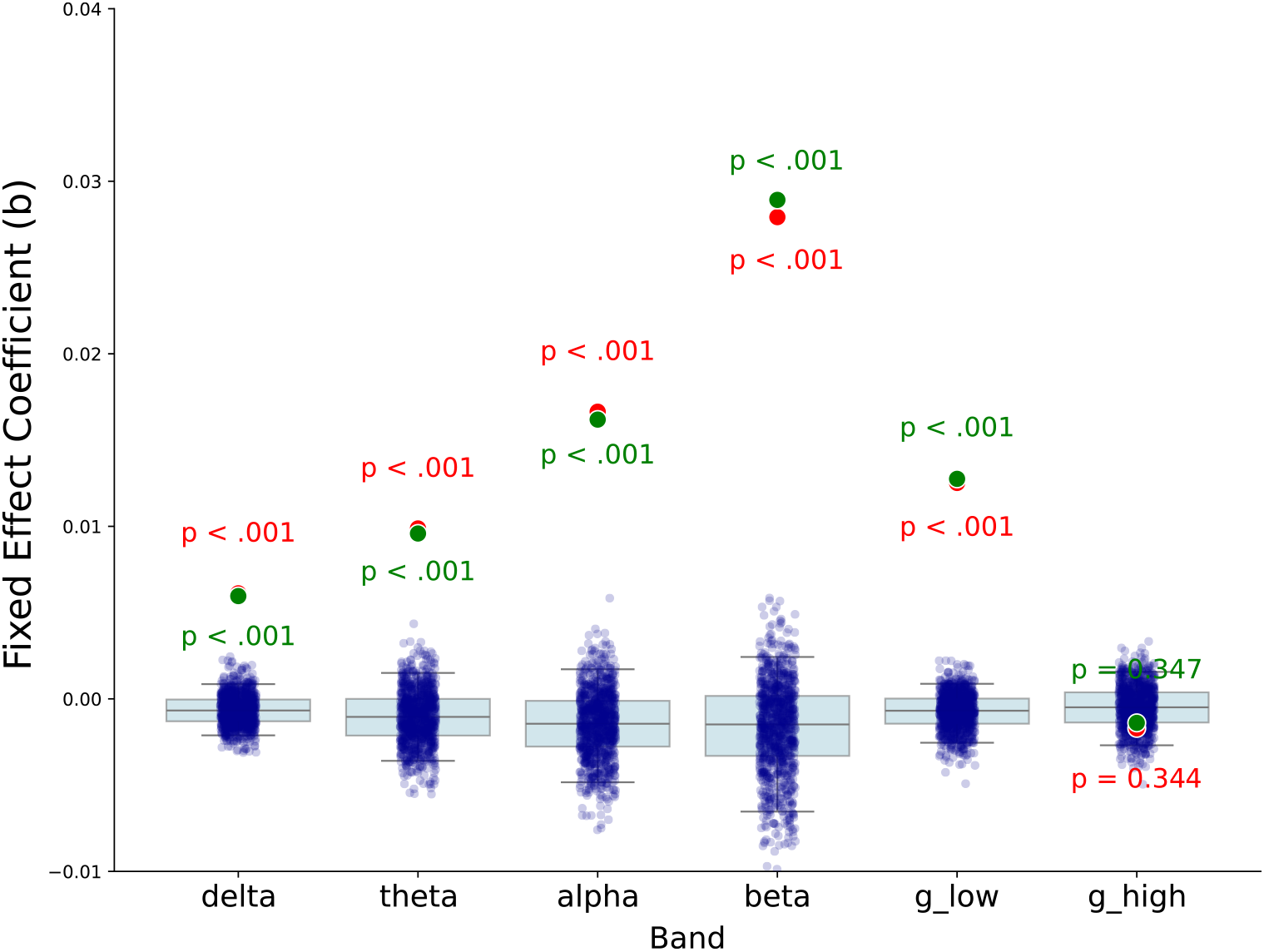
Neurochemical–connectivity alignment is robust to age-related effects. Fixed-effect coefficients (b) for the association between functional connectivity and neurochemical similarity are shown across canonical frequency bands, before (red) and after (green) controlling for age-related structure in the neurochemical similarity matrix. All primary effects (δ, θ, α, β, γ_↓_) remained significant after age correction (p_F DR_ < 0.001), with effect sizes unchanged relative to the main. Background distributions represent spatial autocorrelation-preserving null models.

